# Auditory attention improves scale-invariant neural fidelity to speech across three EEG datasets

**DOI:** 10.64898/2026.08.11.744085

**Authors:** Yu Ding, Jia Zhang

## Abstract

Neural speech tracking is stronger for attended speech, yet its common correlation-based readout is scale invariant, so interpreting this effect only as response gain is incomplete. We tested whether attention improves representational fidelity, defined here as scale-invariant agreement between a speech envelope and its neural reconstruction. A leakage-resistant analysis evaluated held-out trials or story parts in three public electroencephalography datasets (52 participants). Fidelity was Fisher-transformed reconstruction–envelope correlation; projection slope quantified scale-dependent gain. In the spontaneous Auditory Attention Switching Dataset, nine odd-numbered participants were used for discovery and nine even-numbered participants for split-sample validation. Fidelity was higher for attended speech in the validation sample and exceeded 5,000 within-trial circular label shifts. The effect replicated under story-part-disjoint validation in KUL and trial-disjoint validation in DTU. A KUL crossover compared the same clean speech sources in attended and ignored states. Across 4,819 isolated spontaneous switches, fidelity did not differ from baseline before the report but shifted toward the newly reported target 0.25–1 s afterward. Passive keypresses altered nonspecific decoder energy. Gain advantages also occurred in all datasets. Selective attention was therefore evident in the scale-invariant preservation of target dynamics, while gain remained a complementary feature.

## 1. Introduction

Following one talker in a crowded acoustic scene requires the nervous system to preserve behaviorally relevant structure while competing signals reach the ears at the same time. Neural recordings have shown that attended speech dominates cortical population representations and can be reconstructed from electrocorticography, magnetoencephalography, and scalp EEG (Ding & Simon, 2012a; Fuglsang et al., 2017; Mesgarani & Chang, 2012; O’Sullivan et al., 2015). These findings established neural speech tracking as both a mechanistic assay and the basis of auditory-attention decoding. Attention-related alpha lateralization during directed dichotic listening provides a complementary index of spatial selection (Payne et al., 2017), but it does not identify how faithfully the selected speech content is represented. What, exactly, becomes better in the attended representation therefore remains unsettled.

A common explanation is gain: attention increases the magnitude of the target-evoked response or the effective weight assigned to the target. Gain is clearly important in many auditory tasks (Choi et al., 2013; Ding & Simon, 2012b). However, increased response magnitude is not the only possible change. Attention may also improve the temporal or spectrotemporal *fidelity* with which neural activity follows a target. Rimmele and colleagues explicitly described speech tracking as more precise for attended natural speech (Rimmele et al., 2015). Target-enhancement work has likewise noted that increased tracking could reflect sensory gain, more faithful temporal encoding, or both (Orf et al., 2023). Beyond continuous speech, selective attention can sharpen auditory population receptive fields (Lage-Castellanos et al., 2023) and reduce trial-to-trial cortical response variability (Strait et al., 2014). In awake rats, attentional manipulations increased stimulus–response coherence in the deep superior colliculus without corresponding changes in response amplitude, motivating a precision-based account across levels of the auditory system (Ding et al., 2025).

Here, representational fidelity has a deliberately narrow operational meaning: the scale-invariant agreement between a physical target feature and a neural readout of that feature. For continuous speech, we quantify it as the Fisher-transformed correlation between a held-out speech envelope and the corresponding envelope reconstructed from EEG. Multiplying the reconstruction by any positive constant leaves fidelity unchanged. We quantify reconstruction gain separately as the least-squares projection slope. This distinction is measurement-level, not a claim that biological gain and fidelity are causally independent. In a noisy system, larger neural gain can improve effective signal-to-noise ratio and thereby increase correlation. Speech–brain coherence can also change because its periodic and aperiodic components shift with intelligibility, underscoring that a single increase in tracking magnitude is not a unique description of neural coding (Schmidt et al., 2023). The central claim tested here is therefore conservative: selective attention should be observable in a scale-invariant fidelity measure and should generalize beyond a particular stimulus, participant split, or laboratory.

This framing creates three methodological demands. First, evaluation windows from the same trial or story must not be split between training and testing. Random window splits can allow a model to recognize trial-specific fingerprints rather than generalize stimulus–response mapping (Puffay et al., 2023; Rotaru et al., 2024). Second, acoustics must be controlled. A fidelity advantage could otherwise reflect one story, voice, or spatial channel being easier to reconstruct. Third, spontaneous attention reports require explicit treatment of motor activity. The recently released Auditory Attention Switching Dataset (AASD) includes freely initiated switches, button reports, and a passive alternating-keypress condition, which allows potential motor contamination of switch-locked EEG to be assessed (Wang et al., 2026).

We addressed these demands with a common analysis across AASD, the KUL auditory-attention dataset, and the DTU real-world speech dataset (Biesmans et al., 2017; Das et al., 2016; Fuglsang et al., 2018; Wang et al., 2026). AASD was divided deterministically by participant number: odd-numbered participants formed a discovery set used to select the event-to-channel mapping, and even-numbered participants formed a split-sample validation set. This split was not preregistered and is described as internal validation rather than formal confirmation. KUL models left out every occurrence of a story part, including its repetitions. The same clean speech tracks were also compared after attentional status reversed. DTU contributed data from a third laboratory, with different preprocessing and speech materials. We evaluated attended-minus- ignored fidelity during stable attention, its replication in KUL and DTU, and stimulus-specific reconfiguration around spontaneous switches. Reconstruction gain, within-trial circular label shifts, and passive keypress-related decoder energy provided complementary and control analyses.

## 2. Method

### 2.1. Open datasets and inclusion

All analyses used de-identified public datasets and the participant sets provided by their authors; no participant was excluded from stable-attention analyses. Sample sizes were fixed by the available datasets rather than selected through a prospective power analysis. The deterministic AASD split assigned half of its 18 participants to validation, resulting in a validation sample of nine participants; two-sided tests, confidence intervals, within-participant effect sizes, and exact Wilcoxon sensitivity tests are therefore reported. KUL and DTU were retained in full as external replications. Demographic variables were not harmonized across releases, so we report only characteristics documented by the original sources and did not conduct demographic subgroup analyses.

#### AASD

Eighteen normal-hearing Mandarin-speaking adults (18–27 years) listened to two HRTF- spatialized speech streams and switched attention freely, reporting left/right focus with two keys (Wang et al., 2026). We analyzed 60 main-task trials of 60 s per participant (18 h total). The passive ten-trial keypress block was used only for the motor audit. Preprocessed data contained 62 EEG channels at 128 Hz; raw passive EEG was sampled at 500 Hz.

#### KUL

Sixteen participants with audiometrically verified normal hearing (17–30 years; 8 female and 8 male) listened to two Dutch stories under dry dichotic and HRTF conditions (Biesmans et al., 2017; Das et al., 2016). Each participant contributed eight approximately 6-min trials plus twelve repeated approximately 2-min excerpts (about 72 min). Identical story material recurred, making content-grouped validation essential. Analyses used KUL release v2.1 (Das et al., 2026), which corrected the approximately 70-ms EEG–stimulus misalignment documented by the data authors on July 17, 2026.

#### DTU

Eighteen of 19 recorded participants were retained in the public analysis dataset; one was excluded by the original investigators because several trials were missing (Wong et al., 2018). Participants listened to competing Danish speech in simulated anechoic and reverberant rooms. The distributed preprocessed files contained 60 two-talker trials of 50 s, 66 EEG channels, an attended envelope, and an unattended envelope at 64 Hz. Age and sex composition were not reported in the public data article or release materials reviewed for this study.

### 2.2. Signal preprocessing

For AASD, each audio-channel envelope was computed as the magnitude of the analytic signal raised to the 0.6 power, then band-pass filtered at 0.5–8 Hz, standardized within trial, and resampled to 32 Hz. EEG was band-pass filtered at 0.5–8 Hz and resampled from 128 to 32 Hz. For stable-model training, samples before the initial state report and samples within 3 s of any keypress were excluded.

For KUL, clean dry-source audio was resampled to 8 kHz, Hilbert envelopes were compressed with exponent 0.6, filtered at 0.5–7.5 Hz, standardized, and resampled to 16 Hz. EEG was referenced to Cz following the dataset example pipeline, filtered at 0.5–7.5 Hz, and resampled from 128 to 16 Hz.

For DTU, the authors’ auditory-filterbank envelopes (power 0.3) and preprocessed EEG were filtered at 0.5–7.5 Hz and resampled from 64 to 16 Hz. The slightly lower rate for the longer KUL and DTU recordings reduced feature dimensionality while retaining the analyzed band.

### 2.3. Trial-disjoint stimulus reconstruction

For each training fold, EEG channels were standardized using training samples only. A ridge model reconstructed the attended envelope from all EEG channels and lags from 0 through 500 ms. The ridge parameter was α = 100. Sufficient-statistic accumulation (X’X and X’y) avoided concatenating large recordings; every second training origin was used at 16 Hz for KUL and DTU, whereas all 32-Hz samples were used for AASD. Test data were never used to estimate scaling or decoder weights.

AASD used six folds of ten consecutive trials. KUL used four folds defined from the part1–part4 identifier parsed from each audio filename; the held-out fold contained the full trial and all repeated excerpts from that part. DTU used six folds of ten consecutive trials. The reconstructed envelope in a held-out trial was compared with both candidate streams in causal rolling windows. An estimate timestamp equaled the latest EEG sample used by the 0–500 ms decoder lags.

### 2.4. Fidelity, gain, and trial readout

Within each 2-s window, fidelity was defined as the Pearson correlation between the reconstruction and stimulus, followed by Fisher z transformation. The primary margin was attended minus ignored fidelity. Gain was the ordinary least-squares slope from stimulus to reconstruction within the same window, and the gain margin was attended minus ignored slope. Trial readout was correct when a held-out trial’s mean fidelity margin was positive. Window estimates were averaged within trial and participant as appropriate; group tests always used one value per participant.

### 2.5. Discovery and split-sample validation

AASD odd-numbered participants (1,3,…,17; *n*=9) were designated discovery data. Because the public event array contained numeric state codes without a textual channel dictionary, both possible mappings were evaluated in this set. The identity mapping had the larger discovery-set attended-minus-ignored reconstruction margin and was carried forward unchanged to even-numbered participants (2,4,…,18; *n*=9). The 0–500-ms lag interval, 2-s primary window, 3-s key- adjacent exclusion, and α = 100 followed the dataset’s technical-validation settings rather than optimization on the validation participants. No preregistration or contemporaneous time-stamped protocol is claimed for this split-sample analysis.

### 2.6. KUL clean-source-matched crossover

For unrepeated KUL trials 1–8, each story part appeared twice with attention directed to opposite speech sources. Because the held-out part was absent from model training, the same decoder evaluated both presentations. For each clean-source waveform, fidelity when attended was subtracted from fidelity when ignored; the two source differences were averaged across the four parts for each participant. This matches speech content and source envelope, but not necessarily source-to-ear assignment or the complete binaural mixture.

### 2.7. Spontaneous-switch analysis

For temporal resolution, AASD was rerun with a 0.5-s decision window. State transitions after the initial report were switch events. An event was retained only if it and the ±3-s analysis range remained at least 3.5 s from adjacent switches or boundaries. This yielded 4,819 events. Fidelity and gain were signed relative to the newly reported target, interpolated on a 125-ms grid, averaged across events within participant, and then analyzed across participants. The prereport (−1 to −0.25 s), early postreport (0.25–1 s), and late postreport (1.5–2.5 s) means were contrasted against −3 to −2 s baseline with two-sided one-sample tests.

### 2.8. Circular label shifts and passive motor audit

For each AASD participant and main-task trial, the reported state sequence was circularly shifted by a random amount of 5–55 s while fidelity values and the stable-sample mask remained fixed. Five thousand null datasets preserved participant, trial, state occupancy, autocorrelation, and channel-specific reconstruction properties. Validation-set group means formed the null distribution.

For the passive audit, raw 64-channel EEG was referenced to the mastoids, the reference channels removed to match the 62-channel processed data, filtered at 0.5–8 Hz, and resampled to 32 Hz. A decoder fit to all stable main-task data was applied to passive trials. The rolling 0.5-s standard deviation of decoder output indexed nonspecific energy. Isolated passive keypresses and main-task switches were averaged within each validation participant. Because an extreme raw passive value violated normality, the sign-consistent Wilcoxon result is emphasized.

### 2.9. Statistics and reproducibility

Stable-attention contrasts were tested with two-sided one-sample *t* tests against zero (or 0.5 for accuracy), accompanied by exact two-sided Wilcoxon signed-rank tests. Switch contrasts were also two-sided. Two-sided 95% confidence intervals and within-participant Cohen’s *d*_z_ are reported. Shapiro–Wilk diagnostics and exact statistics are provided in the accompanying statistical_results.csv file. No window or event was treated as an independent replicate. Randomization used seed 20260808. For the three fidelity contrasts around spontaneous switches, Holm adjustment gave *p* = .00547 for the early window and *p* = .00200 for the late window; the prereport contrast remained nonsignificant. Unadjusted values are reported with the analysis table, and adjusted values are stated where the contrasts are interpreted.

All processing and statistics were implemented in Python; all final figures were generated with MATLAB R2015a from exported CSV tables. Source scripts, frozen outputs, 600-dpi PNG figures, and vector PDFs accompany this submission.

### 2.10. Use of generative artificial intelligence

Yu Ding conceived and designed the study, prepared the initial analysis code, selected and read the literature, downloaded the public datasets, executed the analyses, interpreted the results, and retained final approval of the manuscript. OpenAI Codex (GPT-5; accessed August 2026) assisted with literature searching, code review and refinement, language editing, reference- format checks, and document formatting. AI-generated suggestions were treated as unverified and were not used as an autonomous source of scientific decisions or interpretation. Yu Ding checked the cited sources, code, numerical outputs, figures, and final text. Selected outputs from the locally executed pipeline were also checked against a frozen machine-readable verification file.

## 3. Results

### 3.1. A scale-invariant operationalization of representational fidelity

For each participant, a regularized linear decoder reconstructed the attended speech envelope from multichannel EEG at lags of 0–500 ms (Figure 1). At every held-out time window, the same reconstruction was compared with both candidate speech envelopes. The primary outcome was

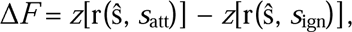

**Figure 1.**
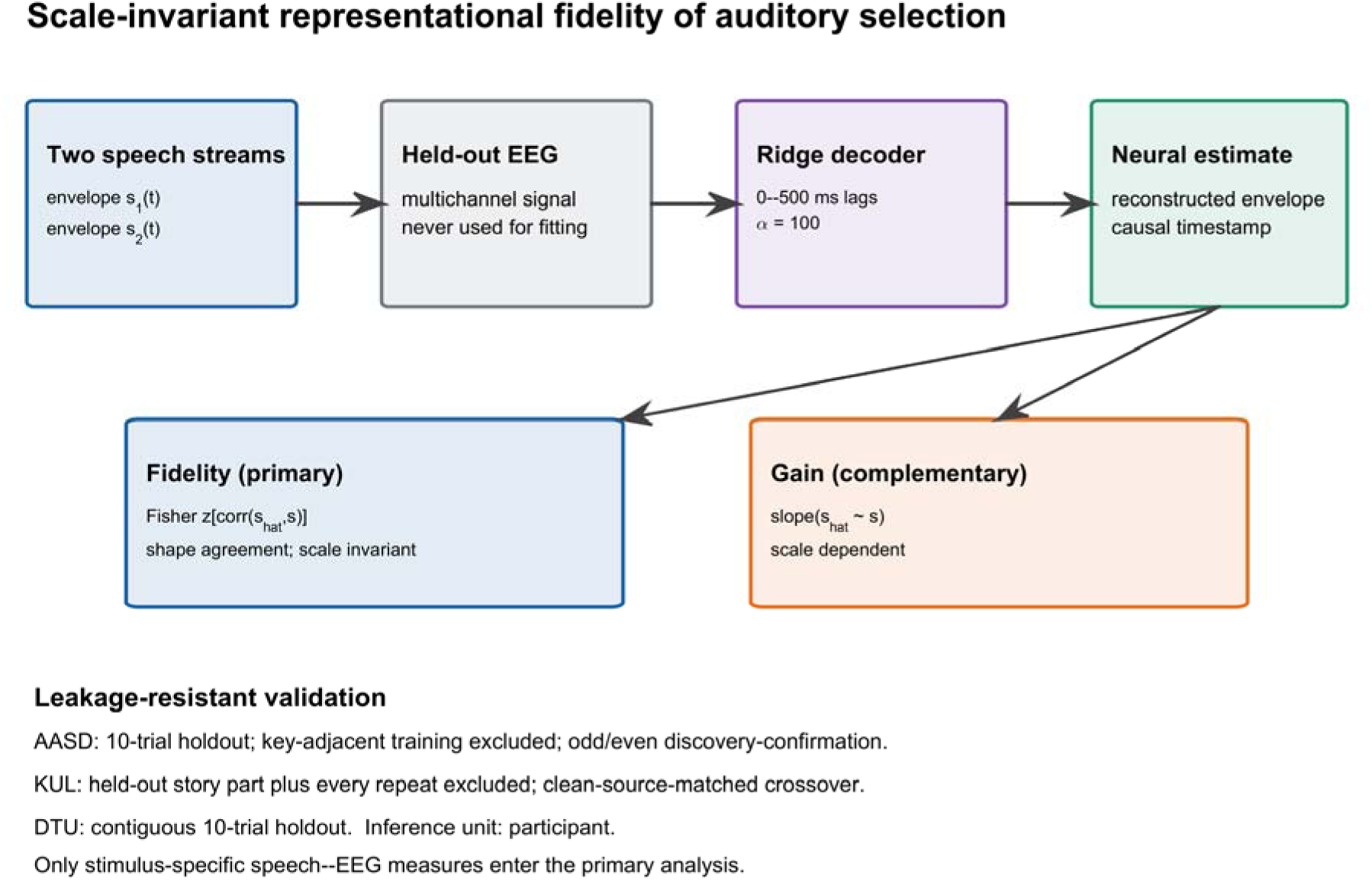
Scale-invariant representational-fidelity framework and validation design. A regularized EEG decoder reconstructs the attended speech envelope from 0–500-ms lags. Fidelity is Fisher-transformed reconstruction–stimulus correlation and is invariant to positive rescaling. Gain is a complementary projection slope. All evaluation is trial/content disjoint; AASD additionally uses an odd/even discovery–validation split and excludes key-adjacent training data.

where *z* is Fisher’s transform, ŝ is the EEG-derived reconstruction, and *s*_att_ and *s*_ign_ are the attended and ignored envelopes. Positive Δ*F* means that the shape of the neural reconstruction more closely follows attended speech. Because correlation is invariant to positive rescaling, this measure does not treat a larger reconstruction as greater fidelity. The complementary gain difference was calculated from regression slopes from *s* to ŝ.

All model evaluation was content-disjoint. AASD used six contiguous 10-trial folds; KUL used four folds that removed every full-length and repeated presentation of the held-out story part; DTU used six contiguous 10-trial folds. Participant—not window, electrode, trial, or switch— was the inferential unit.

### 3.2. Split-sample validation in spontaneous attention switching

The nine odd-numbered AASD participants were used to select the event mapping; the 2-s decision window, 0–500-ms lag range, and ridge penalty (α = 100) followed the dataset’s technical validation rather than optimization on the present validation sample (Wang et al., 2026). Applying this configuration to the nine even-numbered participants yielded a positive fidelity advantage (mean = 0.0474 Fisher-z units, SD = 0.0609, 95% CI [0.0006, 0.0942]; one-sample two-sided *t*_(8)_ = 2.335, *p* = .0478, *d*_z_ = 0.778; exact two-sided Wilcoxon *W* = 6, *p* = .0547; Figure 2a). Six of nine validation participants had positive mean fidelity differences.

**Figure 2.**
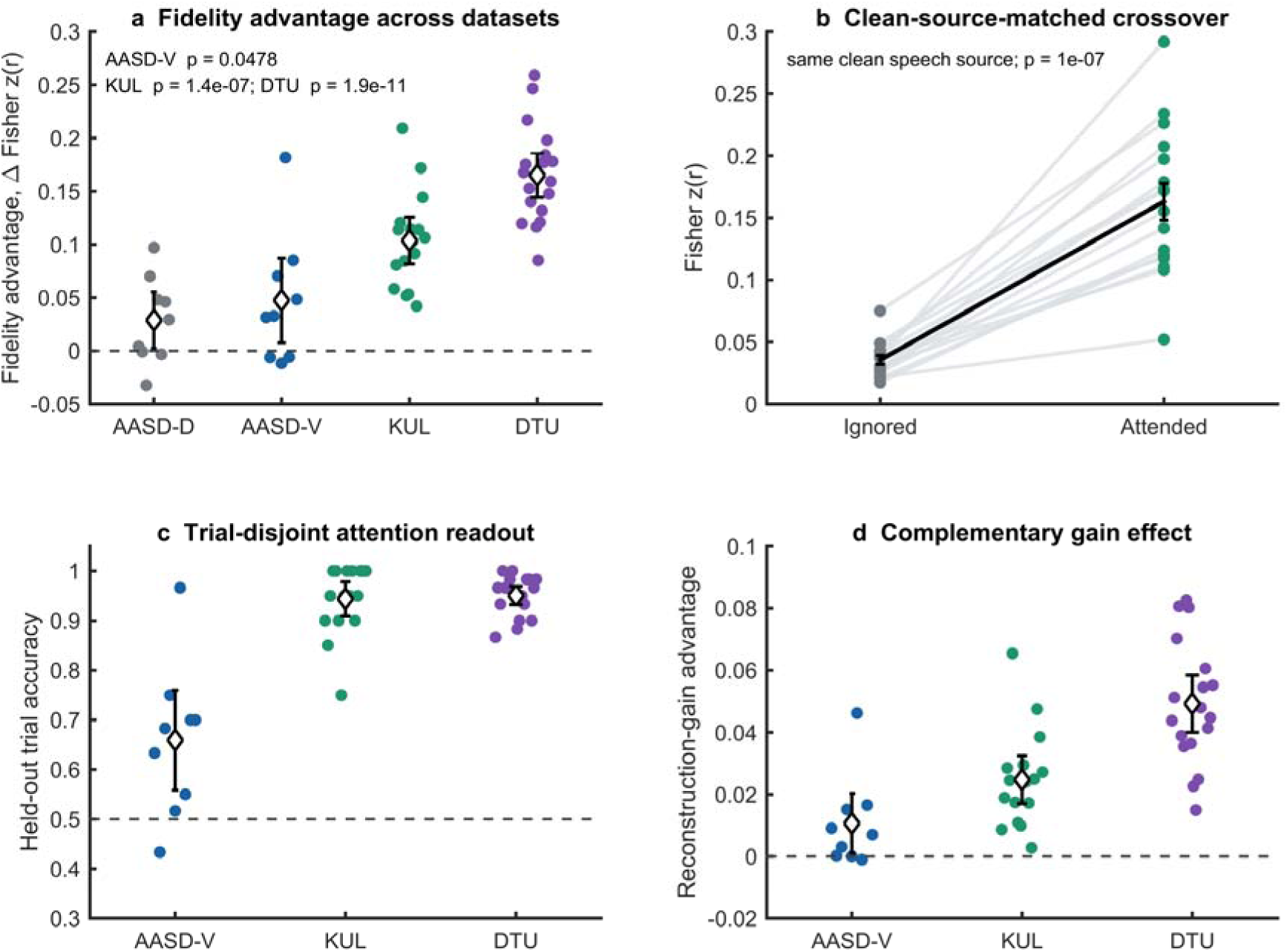
Stable-attention fidelity generalizes across datasets. (a) Participant fidelity margins in AASD discovery, AASD validation, KUL, and DTU. Diamonds and error bars show means and 95% normal-approximation CIs; dots are participants. (b) KUL clean-source-matched crossover: each line joins mean fidelity for the same clean speech source when ignored versus attended; the black line and error bars show the group mean ± SEM. (c) Fully held-out trial readout. (d) Complementary reconstruction-gain margins. Inferential values are participant based.

Aggregating held-out windows within each of the 60 trials yielded a mean participant-level trial accuracy of 65.9% (SD = 15.4%), above 50% chance (*t*_(8)_ = 3.095, two-sided *p* = .0148; Wilcoxon *p* = .0195; Figure 2c). This trial-level measure assesses whether the mean fidelity difference within each fully held-out trial correctly identified the reported attentional state; it is not intended as a window-level decoding benchmark.

The complementary gain difference was positive but uncertain under a two-sided parametric test (mean = 0.0107, *t*_(8)_ = 2.168, *p* = .0620; Wilcoxon *p* = .0195; Figure 2d). The scale-invariant fidelity result therefore does not require a gain-free account of attention. A 0.5-s-window sensitivity analysis retained a positive mean validation-set fidelity difference (mean = 0.0336), although neither two-sided test crossed .05 (*t* test *p* = .110; exact Wilcoxon *p* = .0977).

### 3.3. Story-disjoint replication in KUL and trial-disjoint replication in DTU

The KUL analysis included 16 participants, each contributing approximately 72 min of EEG data. Because this dataset contains repeated excerpts, all occurrences of a story part were assigned to the same test fold. Using the corrected v2.1 files, the fidelity advantage was positive in all 16 participants (mean = 0.1037, SD = 0.0448, 95% CI [0.0799, 0.1276]; *t*_(15)_ = 9.262, two- sided *p* = 1.36×10^-7^, *d*_z_ = 2.32; exact two-sided Wilcoxon *W* = 0, *p* = 3.05×10^-5^). The participant-level held-out trial accuracy was 94.4%. Effects were present separately for dichotic dry stimuli (mean = 0.0964, *p* = 8.46×10^-8^) and HRTF stimuli (mean = 0.1111, *p* = 3.40×10^-7^), and in both unrepeated full-length trials (mean = 0.1275, *p* = 1.00×10^-7^) and repeated excerpts (mean = 0.0879, *p* = 4.12×10^-7^); all subset tests were two-sided.

We also reran the identical pipeline on v2.0 to quantify the effect of the corrected alignment. The mean fidelity margin increased by 0.00164 in v2.1; participant estimates were highly concordant across versions (*r* = .9957), the largest absolute participant change was 0.00949, and the paired version difference was not statistically reliable (*t*_(15)_ = 1.545, two-sided *p* = .143). All 16 v2.1 participant effects remained positive. Primary results and figures use v2.1; v2.0 appears only in this sensitivity comparison.

The DTU analysis included 18 participants, each with 60 50-s trials. Contiguous 10-trial blocks were held out. Again, the fidelity advantage was positive in every participant (mean = 0.1653, SD = 0.0454, 95% CI [0.1427, 0.1878]; *t*_(17)_ = 15.454, two-sided *p* = 1.93×10^-11^, *d*_z_ = 3.64; exact two-sided Wilcoxon *W* = 0, *p* = 7.63×10^-6^). Mean held-out trial accuracy was 95.1%. Reconstruction gain was also higher for attended speech in both KUL (mean = 0.0247, two-sided *p* = 1.43×10^-5^) and DTU (mean = 0.0492, two-sided *p* = 7.81×10^-9^).

Across the validation and external-replication samples, the same directional effect was present in 40 of 43 participants (6/9 AASD validation, 16/16 KUL, and 18/18 DTU). Formal statistical inference was based on continuous participant-level estimates rather than dichotomized sign counts.

### 3.4. Clean-source-matched crossover excludes a speech-content explanation

KUL’s first eight trials contain matched story-part pairs: the same two clean speech sources were presented again with the attended source reversed. Models for a held-out part were trained without any occurrence of that part. We compared each source waveform’s fidelity when it was attended with its fidelity when it was ignored. In v2.1, this clean-source-matched crossover produced a positive difference in all 16 participants (mean = 0.1275, SD = 0.0538, 95% CI [0.0988, 0.1562]; *t*_(15)_ = 9.481, two-sided *p* = 1.00×10^-7^, *d*_z_ = 2.37; exact two-sided Wilcoxon *W* = 0, *p* = 3.05×10^-5^; Figure 2b). Consequently, differences in story identity, talker, envelope modulation, or intrinsic source reconstructability cannot explain the KUL attention effect. Because source-to-ear assignment can reverse between presentations, this control does not equate the complete binaural waveform.

### 3.5. Speech-specific fidelity rapidly reconfigures after spontaneous reports

For exploratory switch dynamics, AASD fidelity was recalculated in 0.5-s causal windows. The timestamp of each estimate was the latest EEG sample used by the decoder. Switches less than 3.5 s from a neighboring switch or trial boundary were excluded, leaving 4,819 events. Events were averaged within participant before group inference.

Fidelity was expressed relative to the newly reported target. It was negative before the keypress, suggesting that the previous target remained better represented, and became positive approximately 0.5 s after the report (Figure 3a). Compared with a −3 to −2 s baseline, the −1 to −0.25 s pre-report period did not change (mean contrast = −0.0036, two-sided *t*_(17)_ = −0.373, *p* = .714; Wilcoxon *p* = .702). Thus, these data do not support a pre-switch collapse in fidelity. The new-minus-old fidelity increased at 0.25–1 s (mean contrast = 0.0528, *t*_(17)_ = 3.502, two-sided *p* = .00273; Wilcoxon *p* = .00475; Holm-adjusted *p* = .00547) and remained increased at 1.5–2.5 s (mean = 0.0681, *t*_(17)_ = 4.152, *p* = .000667; Holm-adjusted *p* = .00200). Gain showed the same general post-report direction but a smaller standardized early effect (*d*_z_ = 0.57 versus 0.83 for fidelity; Figure 3b, Figure 3c).

**Figure 3.**
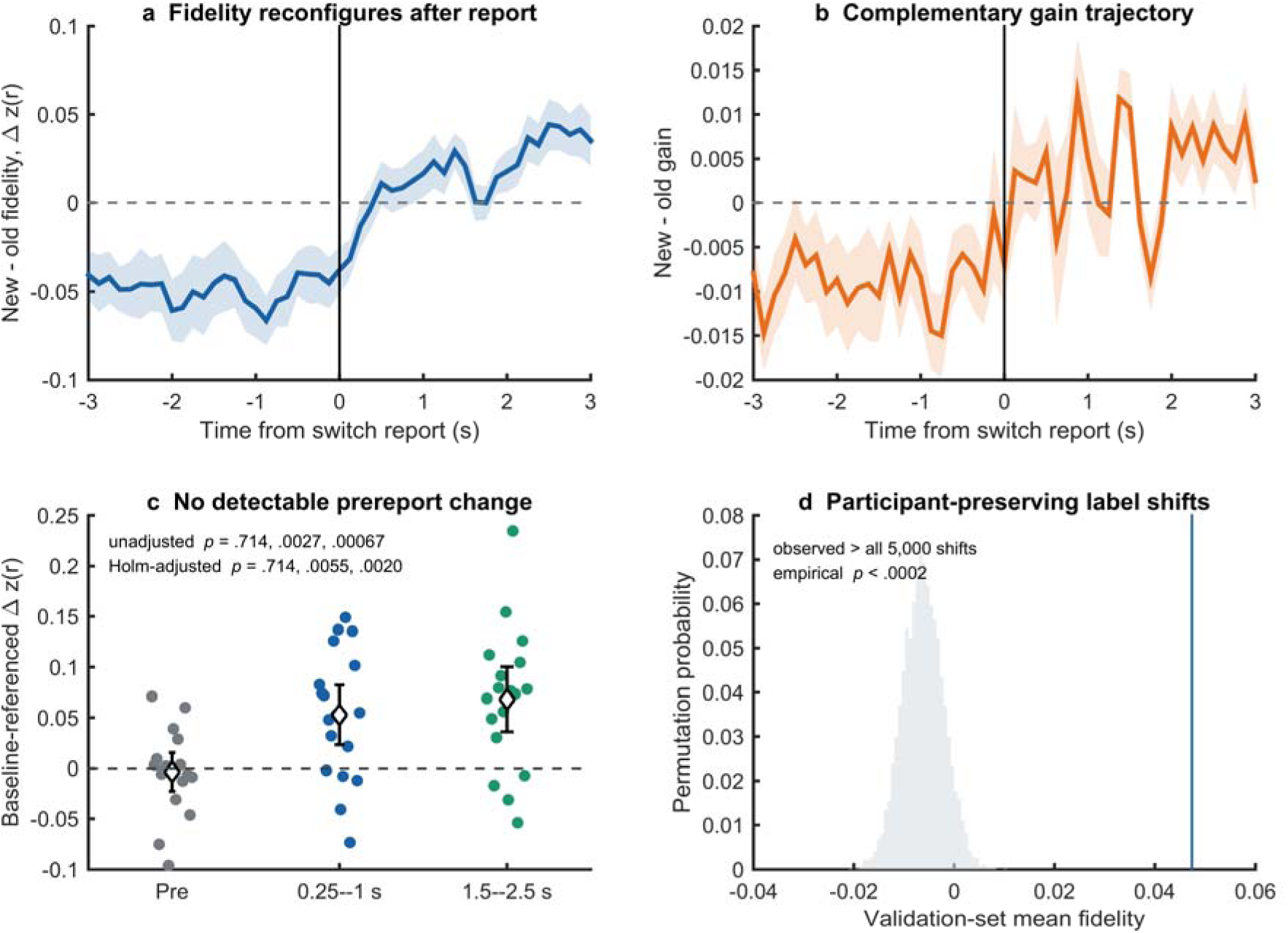
Speech-specific fidelity during spontaneous AASD switches. (a,b) New-target minus old-target fidelity and gain in 0.5-s causal windows; curves are participant means ± SEM after averaging isolated events within participant. (c) Participant fidelity contrasts relative to the −3 to −2 s baseline; labels give unadjusted and Holm-adjusted two-sided *p* values for the prereport, early postreport, and late postreport contrasts, respectively. (d) Validation-set observed fidelity compared with 5,000 participant-preserving within-trial circular label shifts.

These timings describe the estimator and the behavioral report jointly; they are not claimed as absolute neural latencies. The 0.5-s correlation window and the decoder’s response lags necessarily smooth the underlying transition. The key result is the ordering: no detectable fidelity loss preceded self-report, whereas speech-specific fidelity shifted promptly afterward.

### 3.6. Temporal specificity and motor audit

The AASD paper cautions that switch labels are button-locked (Wang et al., 2026). We therefore trained only on samples at least 3 s from any keypress and evaluated only held-out trials. In the even-numbered validation set, the observed mean fidelity advantage exceeded every one of 5,000 participant-preserving, within-trial circular label shifts by at least 5 s (empirical *p* = 1/5001 < .0002; Figure 3d). Static channel preference, state occupancy, and slowly varying trial structure therefore did not reproduce the observed temporal alignment.

We also applied a decoder trained on the main task to raw EEG from the passive block, in which participants alternated the same keys without changing auditory attention. Nonspecific decoder energy changed around passive keypresses in all nine validation participants (exact two-sided Wilcoxon *W* = 0, *p* = .00391; Supporting Figure S1). One participant had an extreme raw-block value, so the sign-based result is emphasized. Generic EEG magnitude is therefore motor-sensitive and is not interpreted as a specific measure of attention. The primary endpoint additionally requires moment-by-moment agreement with the correct held-out speech envelope; the training, trial-holdout, and temporal-shift controls address a nonspecific motor explanation.

## 4. Discussion

Across three public EEG datasets, attended speech was reconstructed with higher scale-invariant fidelity than ignored speech. The evidence included an internal discovery/validation split in spontaneous attention, leave-story-part-out validation, an external dataset from a third laboratory, varied languages and stimulus conditions, and a clean-source-matched attention crossover. During spontaneous switches, fidelity moved from the previous to the newly reported target without a detectable pre-report change. Selective auditory attention was thus measurable in how faithfully neural activity preserved target-speech dynamics after readout scale was removed.

### 4.1. Fidelity is a useful level of description, not a renamed amplitude

Correlation-based speech tracking is not new (Ding & Simon, 2012a; Fuglsang et al., 2017; Mesgarani & Chang, 2012; O’Sullivan et al., 2015). The advance is to make its scale invariance the hypothesis, to contrast it explicitly with a scale-dependent gain estimate, and to subject it to controls targeted at alternative explanations. Multiplying a reconstruction by a constant changes the slope but not correlation. The clean-source-matched crossover further suggests that higher fidelity is associated with attentional status rather than with a speech source that happens to be easier to encode. These features connect human natural-speech results to work showing attention-related alpha modulation, receptive-field sharpening, reduced response variability, and enhanced stimulus–response coherence (Ding et al., 2025; Lage-Castellanos et al., 2023; Rimmele et al., 2015; Strait et al., 2014).

The present data do not demonstrate that biological gain and precision are independent mechanisms. Gain advantages were robust in all three datasets, and raising the target response relative to background neural noise can improve correlation. A stronger causal claim would require repeated neural responses to identical mixtures under controlled response-amplitude manipulations or a generative model with separately identifiable gain and noise terms. Our conclusion is therefore primarily measurement-level: an attention effect remains detectable after removing the scale of the neural readout, indicating that response magnitude alone is insufficient to describe the effect.

### 4.2. Dynamic selection in spontaneous and cued switching

Recent cued-switch work reported that engagement with a new stream can begin before disengagement from the old stream is complete, creating a period of simultaneous cortical representation (Carta et al., 2026). In AASD, the event marker was a self-initiated report rather than an instruction cue, meaning that the two studies anchor their analyses to different points in the switching process. We found no reliable change before the report and a rapid sign reversal afterward. The public labels do not reveal when the intention to switch arose or whether the keypress occurred before, during, or just after covert reorientation. Future experiments should combine continuous confidence or intention reports with stimulus-specific neural tracking.

### 4.3. Likely level in the auditory hierarchy

Scalp envelope reconstruction predominantly reflects cortical activity. Recent simultaneous measurements found robust cortical attentional modulation from approximately 38–271 ms but no corresponding auditory-nerve or brainstem modulation for natural speech (Stoll et al., 2025). These findings constrain the interpretation of the present effect: we have established a scalp- accessible cortical signature, not evidence that the same effect must occur in the human brainstem. The prior deep-superior-colliculus coherence result (Ding et al., 2025) makes hierarchical tests compelling, but species, task, and measurement differences preclude direct equivalence. Simultaneous cortical and subcortical recordings with the same fidelity estimator are the appropriate next step.

### 4.4. Generalization and safeguards against decoding bias

Direct classifiers can exploit eye position, trial identity, or recording drift instead of stimulus-specific neural encoding (Gehmacher et al., 2024; Puffay et al., 2023; Rotaru et al., 2024). Our models cannot obtain a positive score merely by recognizing a held-out trial’s attention label: they must reconstruct time-varying speech content and correlate more strongly with the correct envelope. KUL’s exact repeated content never crossed the train/test boundary, and the crossover held acoustics constant. Nonetheless, ocular activity itself can track attended speech (Gehmacher et al., 2024). Without simultaneous eye tracking in all datasets, a shared oculomotor contribution to the scalp signal cannot be completely excluded. This limitation concerns the biological source of the stimulus-locked signal, not the existence of the attention-dependent fidelity relationship.

### 4.5. Limitations and future experiments

First, representational fidelity was restricted to the slow speech envelope (0.5–8 Hz or 0.5–7.5 Hz). The conceptual framework motivating this work distinguishes envelope and temporal fine structure. Scalp EEG at the present sampling and SNR cannot provide a strong test of fine- structure fidelity; MEG, frequency-following responses, or intracranial recordings are needed. Second, public datasets differed in language, referencing, envelope extraction, sample rate, and trial structure. This heterogeneity strengthens conceptual generalization but prevents direct comparison of raw effect magnitudes. Third, AASD switches were self-reported by keypress and lacked an independent behavioral marker of switch completion. Fourth, linear reconstruction captures only a subset of auditory coding; periodic and aperiodic tracking components, context-dependent responses, and nonlinear representations may contain additional attention effects (Schmidt et al., 2023; Wikman et al., 2025). Fifth, source localization was not attempted, so this study addresses expression fidelity at the scalp rather than transmission fidelity between named brain regions.

A prospective experiment should repeat identical binaural mixtures under crossed attention while recording high-density EEG/MEG, brainstem responses, eye tracking, and trial-wise comprehension. It should independently manipulate stimulus level to identify gain, measure trial-to-trial residual variability after optimal scaling to identify precision, and use directed cross-validated models to test whether precision changes propagate from auditory cortex to frontoparietal control regions. Such a design would directly connect expression precision, transmission precision, and behavior.

## Open practices statement

### Data availability

AASD is available at https://doi.org/10.5281/zenodo.17413336. KUL v2.1, used for the primary KUL results, is available at https://doi.org/10.5281/zenodo.21415059 (Das et al., 2026). KUL v2.0 (https://doi.org/10.5281/zenodo.4004271) was retained only for the version-sensitivity comparison. DTU is available at https://doi.org/10.5281/zenodo.1199011. No raw participant data are redistributed with this submission.

## Code availability

All analysis and figure-generation code, configuration templates, a step-by-step reproduction manual, and nonidentifying derived verification results accompany the submission as a review archive. The code will be deposited in a permanent public repository no later than acceptance, and the final repository link will replace this sentence before production.

## Ethics statement

This study reanalyzed de-identified public data, recruited no new participants, and made no attempt at reidentification. For AASD, the Southern University of Science and Technology Institutional Ethical Review Board approved the protocol (No. 2022DZX003); participants gave written informed consent, including consent for public sharing of anonymized data (Wang et al., 2026). The KUL protocol and consent form were approved by the KU Leuven ethics committee, and participants or their legal guardians provided written informed consent (Biesmans et al., 2017). The DTU protocol was approved by the Science Ethics Committee for the Capital Region of Denmark, and participants provided written informed consent in accordance with the Declaration of Helsinki (Wong et al., 2018). The author should confirm whether the home institution requires a formal exemption or determination for this public-data secondary analysis before submission.

## Funding

This work was supported by grants from the National Natural Science Foundation of China (NSFC: 32500941). This work was also supported by the Youth Foundation for Humanities and Social Sciences Fund of Ministry of Education of China (grant number 25YJC740070).

## Conflict of interest

The authors have no conflicts of interest to declare.

## Acknowledgments

The author thanks the investigators who made the AASD, KUL, and DTU datasets publicly available. No individuals are named here without their permission.

## Notes

### Competing Interest Statement

The authors have declared no competing interest.

