## Supplemental Figure S1 for "Auditory attention improves scale-invariant neural fidelity to speech across three EEG datasets"

### Supporting Information

**Supporting Figure S1. Passive motor audit of nonspecific decoder energy.** The same main-task decoder responds in energy around both attention-switch reports and passive keypresses. This demonstrates why generic decoder magnitude is not interpreted as attention. Fidelity additionally requires alignment with held-out speech content, training excludes key-adjacent samples, and true alignment exceeds circular shifts.

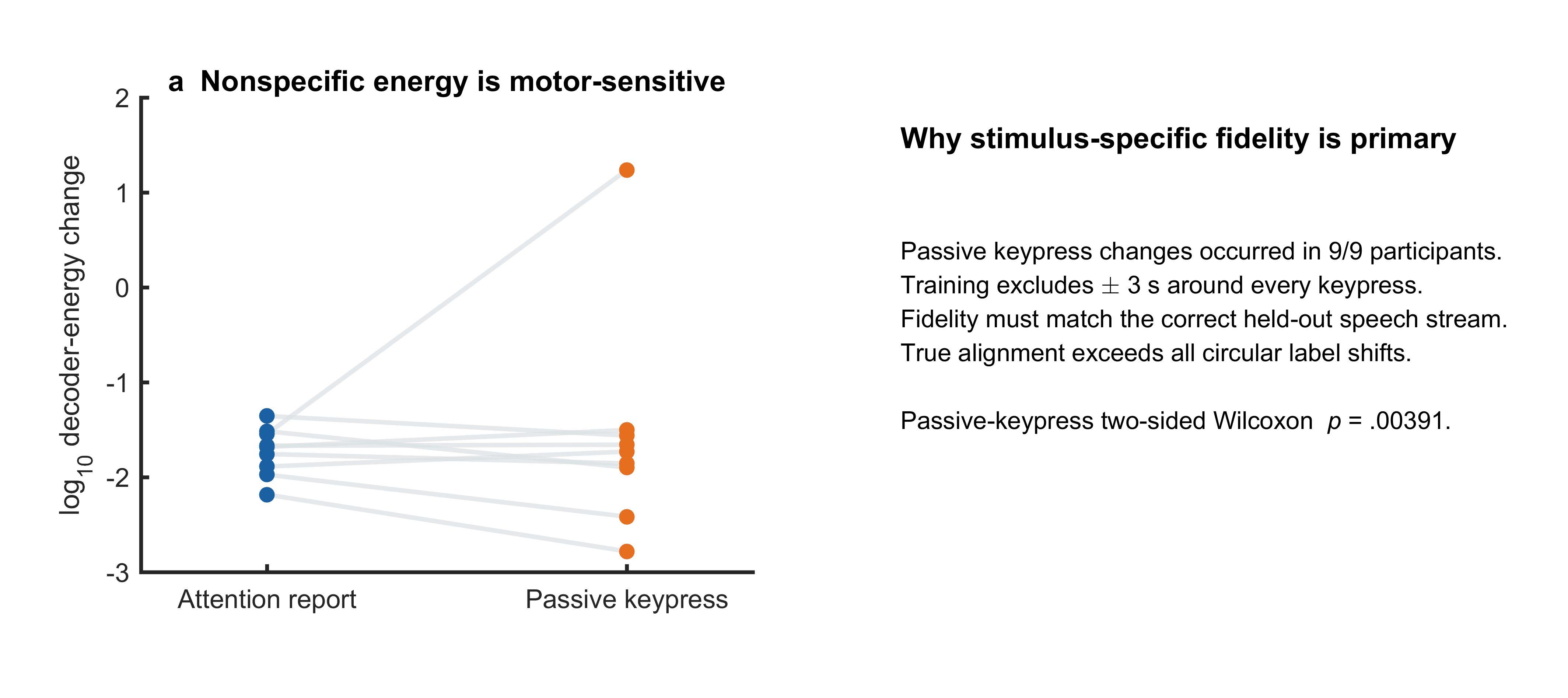
